# Complete genome sequences of nine isolates from microplastics in the Bow River, Calgary, Canada

**DOI:** 10.1101/2024.10.19.619244

**Authors:** Kira L Goff, Sneha Suresh, Jonas M Stadfeld, Srijak Bhatnagar

**Author notes:** Address correspondence to Srijak Bhatnagar. Kira L Goff and Sneha Suresh contributed equally to this work. Author order was determined based on seniority.

## Abstract

We present the complete genome sequences of nine bacterial strains isolated from microplastics from water or sediments of the Bow River in Calgary, Alberta. These isolates provide insight into the freshwater microplastic microbiome and their plastic biodegradation potential.

## Introduction

Microplastics are pervasive contaminants in environments throughout the globe, serving as hosts for microbial communities distinct from those in the surrounding environment (1). Here, we present genomes of bacteria isolated from microplastics in the Bow River, after the confluence with Nose Creek and Elbow River, within Calgary, Canada’s fourth largest city.

Microplastics were isolated from water (40 L) and sediment samples (0.5 Kg) (51.04344 °N, 114.01502 °W) through 20µm filtration or CaCl_2_ density separation (1.4 Kg/L), respectively, followed by Nile red staining (2). Microplastics were rinsed twice with 1x PBS buffer and inoculated in either liquid LB (3) or R2A media (4). Upon visible growth, a 10^−4^ dilution of the liquid culture was spread-plated on the respective solid media, followed by 4-5 rounds of streaking to achieve isolation. One colony from each isolate was grown in respective liquid media for 48 hours and used for high molecular weight DNA extraction using the Lucigen MasterPure Complete DNA & RNA Purification Kit (Mandel Scientific) using the manufacturer supplied protocol for Gram-positive bacteria. DNA was quantified using the Qubit HS dsDNA kit (Thermo Fisher Scientific Inc.) and assessed for purity with NanoDrop (Thermo Fisher Scientific Inc.). DNA was sequenced on an R10.4.1 Nanopore MinION flowcell using Native Barcoding Kit 24 (SQK-NBD114.24, Oxford Nanopore) and basecalled using Dorado (v7.3.11, e8.2_400bps_sup@v5.0.0 model).

Technical sequences and low-quality reads (≤1000bp, Q ≤10) were removed by Porechop ABI (v0.50, ABI mode) (5) and Chopper (v0.80) (6), respectively. Genomes were assembled with Flye (v2.9.3; nano-hq mode) (7) and polished with Medaka (v1.11.3). Genome completeness and contamination were calculated using CheckM2 (v1.0.1) (8), and CRISPR regions were identified using CRISPR-CasFinder (subtype clustering model) (9). Genomes with ≥99.90% ANI were dereplicated using OrthoANIu (10) and annotated using NCBI Prokaryotic Genome Annotation Pipeline (PGAP, version 6.7) (11). Taxonomy was assigned using GTDB-Tk (v2.4.0 with release 220 database) (12).

The nine isolate genomes with fully circularized chromosomes ranged in size from 3,939,831 to 7,333,835 base pairs, encoding 3,610 to 6,553 coding sequences (CDS) and 72 to 118 tRNAs. Extrachromosomal elements (ECEs) were present in four genomes, with the RS8 genome containing six ECEs. Among the isolates, *Lysinibacillus, Peribacillus*, and *Paenibacillus* are Gram-positive, while *Pseudomonas, Stenotrophomonas*, and *Acinetobacter* are Gram-negative. Four genomes had hits for experimentally verified plastic-degrading genes (identity > 80%) (13, 14). Isolate LW8 has genes for PHA depolymerase (biodegradation of poly(3-hydroxybutyrate-co-3-hydroxyvalerate, polycaprolactone, polyethersulfone, polyhydroxyalkanoate, and polylactic acid) and esterase (polyethylene terephthalate). RS7 contained oxidoreductase Oxr1 (biodegradation of polyurethane and polybutylene adipate terephthalate), and RW6 had a hit for phthalate dioxygenase reductase (phthalate biodegradation). RW9 contained PHA and PHB depolymerase genes involved in the biodegradation of polyhydroxyalkanoate and polyhydroxybutyrate, respectively. Furthermore, two genomes of these genomes (LW8 and RW6) had alkane degradation marker genes (15). Alkane hydroxylases, such as AlkB are known to degrade hydrocarbon-based plastics (16, 17).

## Data Availability

Genomes have been deposited in NCBI Genbank under the Bioproject acc# PRJNA1145385.

## Acknowledgments

We want to acknowledge funding from the Research Incentive Grant and Academic Research Fund of Athabasca University to SB and support from the Natural Sciences and Engineering Research Council of Canada as an Undergraduate Student Research Award to SS and Canada Graduate Scholarship-Master’s program to JMS. We would also like to thank Ruchita Solanki for helping with the preparation of the Nanopore library.

---

**Table 1:** Summary of Genomes.

| Isolate ID | Classification | Genome Size (bp) | GC % | Number of Chromosomes + ECE* | Size of Chromosome + ECE (bp) | CDS | tRNA | CRISPR Repeats | Plastic Degradation Genes | Hydrocarbon Degradation Genes | GenBank Accession |
| --- | --- | --- | --- | --- | --- | --- | --- | --- | --- | --- | --- |
| LS1 | *Lysinibacillus sp.* | 4,761,317 | 37.6 | 1 + 0 | 4,761,317 + 0 | 4,602 | 107 | 1 |  |  | CP167119 |
| LW8 | *Pseudomonas sp.* | 5,952,450 | 60.0 | 1 + 1 | 5,902,350 + 50,100 | 5,277 | 70 | 2 | PHA depolymerase, lipase, esterase | LadA | CP168137 - CP168138 |
| RS5 | *Lysinibacillus sp.* | 5,165,228 | 37.1 | 1 + 0 | 5,165,228 + 0 | 4,900 | 109 | 14 |  |  | CP169418 |
| RS7 | *Peribacillus sp.* | 5,809,518 | 39.8 | 1 + 2 | 5,599,460 + 192,986; 17,072 | 5,508 | 83 | 6 | Oxr1 Oxidoreductase |  | CP168134 - CP168136 |
| RS8 | *Paenibacillus sp.* | 7,333,835 | 43.6 | 1 + 6 | 7,098,594 + 190,342; 19,443; 15,490; 5,758; 2,512; 1,709 | 6,547 | 88 | 8 |  |  | CP168128 - CP168133 |
| RS10 | *Lysinibacillus sp.* | 5,165,155 | 37.1 | 1 + 0 | 5,165,155 + 0 | 4,995 | 109 | 14 |  |  | CP167122 |
| RS11 | *Lysinibacillus sp.* | 4,918,501 | 37.0 | 1 + 4 | 4,867,375 + 19,870; 14,416; 10,960; 5,881 | 4,779 | 119 | 13 |  |  | CP168123 - CP168127 |
| RW6 | *Acinetobacter sp.* | 3,939,831 | 38.8 | 1 + 0 | 3,939,831 + 0 | 3,651 | 73 | 0 | Phthalate dioxygenase reductase | AlkB (2), LadA, AlmA | CP167121 |
| RW9 | *Stenotrophomonas sp.* | 4,537,953 | 66.1 | 1 + 0 | 4,537,953 + 0 | 4,049 | 72 | 8 | PHA<br>depolymerase,<br>PHB<br>depolymerase |  | CP167120 |

71 \* Extrachromosomal Elements

